# Global assessment of interventions for mitigation of amphibian fungal disease is dominated by geoeconomic trends and antifungal success

**DOI:** 10.1101/2025.07.03.662529

**Authors:** Luke E. B. Goodyear, Valentina Concha-Toro, Daniel Pincheira-Donoso

## Abstract

Globalisation networks—a rapidly expanding factor behind Anthropocene environmental erosion—facilitate the accelerating spread of invasive species and diseases globally. Among wildlife infectious diseases, chytridiomycosis (caused by the fungal pathogen *Batrachochytrium dendrobatidis*, or *Bd*) has been widely implicated with the alarming global decline of amphibians— nature’s most endangered organisms. Over the last few decades, a range of interventions have been tested to mitigate the annihilating impacts of *Bd* on amphibian communities, with high disparity in success. However, no integrative quantitative assessment of the effectiveness of interventions exists, thus hindering a broader understanding about where actions and further research should be directed. Using a novel, globally comprehensive database of interventions, we show that antifungal treatments of itraconazole applied to individual tadpoles seem offer the best results compared to other interventions. Further, trials performed *ex situ*, predominantly applied in the Global North, may severely bias the ability to assess the effectiveness of treatments applied in more biodiverse areas, across species that have been most severely affected. Our study is a call to continue experimentation on promising novel methods and to expand the geographic and taxonomic deployment of interventions to approach a robust evidence-based understanding of the effectiveness of treatments.

## INTRODUCTION

A prevailing consequence of industrial activity is the rapid and global spread of invasive species and infectious diseases facilitated by the expansion of globalization corridors (1–5). While most scientific and management efforts have concentrated around the mitigation of the detrimental socioeconomic impacts resulting from emerging infectious diseases, the global spread of wildlife infectious pathogens has gained significant attention in recent years, due to their role in driving rapid and irreversible erosions and extinctions of biodiversity (6,7). Over the last few decades, numerous impactful wildlife diseases have been identified as drivers of animal biodiversity loss (8–10). However, the magnitude of the fungal infectious disease chytridiomycosis as a driver of amphibian declines globally remains unparalleled (11–14). The rapid spread of the causative pathogen, *Batrachochytrium dendrobatidis* (*Bd*), has eroded communities of amphibians across most lineages and climatic regions worldwide, leading this panzootic to be widely considered the most destructive wildlife disease to-date (14).

In addition to *Bd*, a newer form of chytridiomycosis caused by a sister pathogen, *Batrachochytrium salamandrivorans*, has raised further concerns about the disease-driven decline of amphibians (15,16). Therefore, the environmental impact of chytridiomycosis has become one of the major concerns of the Anthropocene extinction crisis (12,14), with a rapidly growing interest in developing strategies for its management and mitigation. Until recently, the deployment of interventions (e.g. drugs) designed to mitigate the impacts of infectious diseases on animals has been motivated nearly exclusively by the need to manage costs for sectors of socioeconomic relevance, such as the agricultural industry or public health. For example, diseases such as the viral rinderpest (17,18) and rabies (19–21) have been targeted with successful intervention programmes. However, with increasing active commitment to mitigate drivers of biodiversity erosion, the design and implementation of intervention programs to manage the impacts of wildlife- specific diseases has been on the rise, with the amphibian disease chytridiomycosis at the forefront of this research challenge.

A significantly broad range of intervention approaches have been envisaged and trialled to manage the global-scale defaunation of amphibian biodiversity both in experimental and natural settings. Such interventions include strategies as diverse as alterations of the physical (22–24) or climatic (e.g. (25–27)) environmental conditions surrounding infected species; augmenting the microbiome with commensal bacteria (e.g. (28,29) or inoculations (e.g. (30,31)); removal of individuals from populations (23,32): and, direct individual application of chemical treatments, such as anti-fungals, herbicides, or insecticides, to eliminate infections (e.g. (33–36)). For decades, the results of these intervention methods from across the world have been widely reported in the literature. However, no integrative analyses have investigated the patterns of success from this array of techniques. Therefore, which treatments are likely to be the best choices remains an open question.

Here, we implement the first comprehensive global-scale analysis of the differential success of interventions from the quantitative assessment of 315 treatment interventions carried out over a quarter of a century (between 2000-2023). We aim to elucidate large-scale patterns of: (1) the types of treatment being trialled; (2) the geographical spread of trials; (3) the range of amphibian species and life stages used in trials; and, (4) the breakdown of studies that have been trialled *in situ/ex situ*, and on the habitat or on the individual amphibian, to ultimately establish for the first time what types of intervention strategies are more successful. By including the scalability and longevity of treatments in our success calculation, we intrinsically consider the possibility of large-scale implementation, which will be of practical import in conservation management decision-making. Collectively, our study will enable the identification of gaps in our understanding of interventions, to then provide recommendations for future research to mitigate the global declines of amphibians due to chytridiomycosis.

## RESULTS

Our results reveal that interventions are predominantly applied *ex situ* (283; 90%), and focused on testing single chemical treatment options, with an emphasis on itraconazole (213; 68%). Only 46 experiments (15%) tested combinations of treatment types in the same experiment. In the majority of cases (224; 71%), the implementation was treatment-based rather than preventive, and a nearly equal number of interventions were trialled on individuals (147; 47%) as applied to the habitat (155; 49%), with only 4% being a combination of both. Interestingly, the majority of the interventions— both *ex situ* and *in situ—*have been implemented across high income countries, with the exception of one intervention in Montserrat, seven populations from South Africa (Figure 1), and 37 interventions with no information on population origin and/or experiment location.

**Figure 1.**
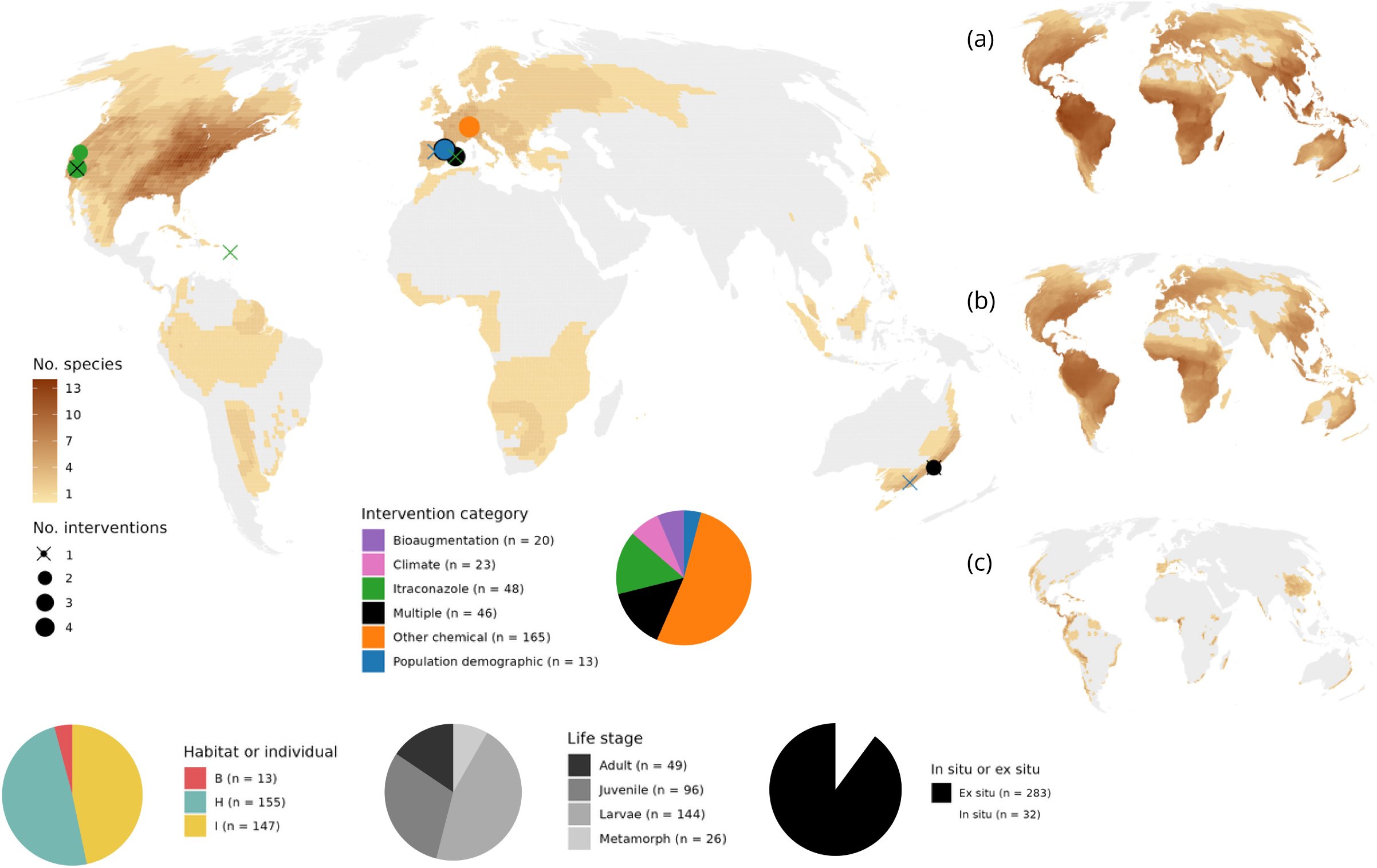
Map showing locations of in situ interventions as points, with the colour of the point indicating the category of intervention and the size of the point indicating number of interventions in that area, where a cross indicates a single intervention. Heat map indicates species ranges for the 61 species that have been used in interventions. Interventions have predominantly been performed in temperate regions using temperate amphibians. Maps down right-hand-side were generated using data from Goodyear et al. 2025 and show the geographical distribution of (a) amphibian biodiversity; (b) *Bd* positive species; and (c) threatened species according to International Union on the Conservation of Nature (IUCN) Red List categories (Vulnerable/Endangered/Critically Endangered).

### Intervention success

We observed a significant degree of differential success among intervention categories, varying across the entire 0–1 score range, with a mean of 0.30 and a median of 0.125 (Supplementary Figure S1a), however all intervention categories received a score of 0 for at least one trial. In total, 102 (32%) interventions had a success score of 0, meaning they either failed to be effective (through a lack of reduction in *Bd* load or prevalence, or failed to increase survivability) or had extreme side effects. Only 15 (5%) interventions scored for longevity, whereas 241 (77%) interventions scored for scalability, however, this is likely because very few interventions had data available for longevity scores, and any data deficient trial was graded with zero for longevity as default. A total of 11 (3%) interventions scored for both, but only five of these had non-zero success scores. Two of these received an overall success score of 1 (the only two interventions to achieve this score) and three scored 0.25. Figure 2 shows the breakdown of specific treatment type within each intervention category, ordered by most frequently trialled, and split by binned success scores, with those involved in multiple treatment interventions being patterned. Success varied across treatment type (Figure 3) with the chemical disinfectant Virkon S being the most promising, particularly as part of a combination treatment plan. Multiple treatments included various combinations, including the two top scorers (Figure 4).

**Figure 2.**
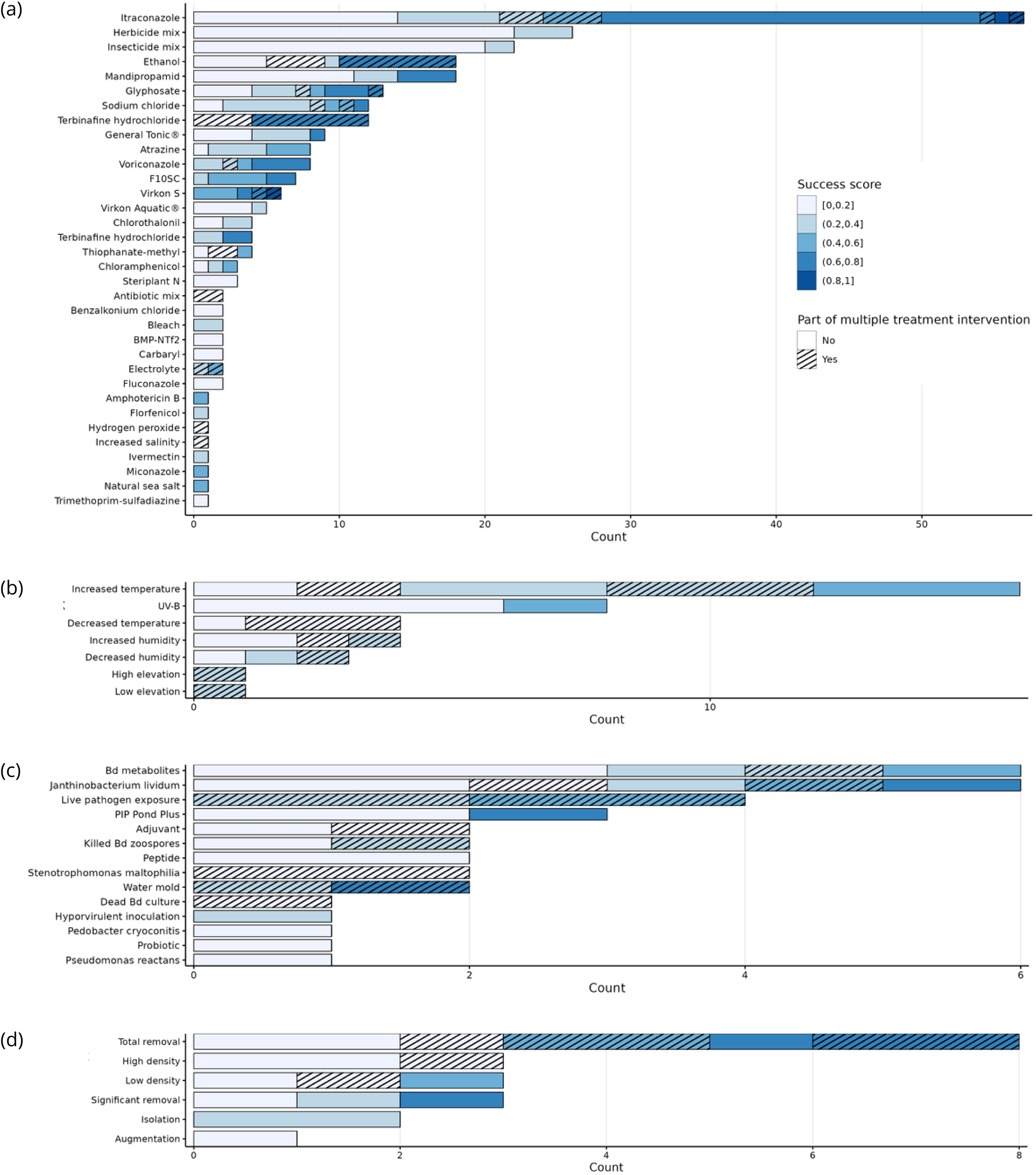
Plot showing breakdown of counts of specific treatments per binned success score. Grouped according to intervention category: (a) chemical treatments; (b) climate treatments; (c) bioaugmentation treatments; (d) population demographic treatments. Colour represents success score and striped pattern marks interventions that were part of a trial with multiple treatments types.

**Figure 3.**
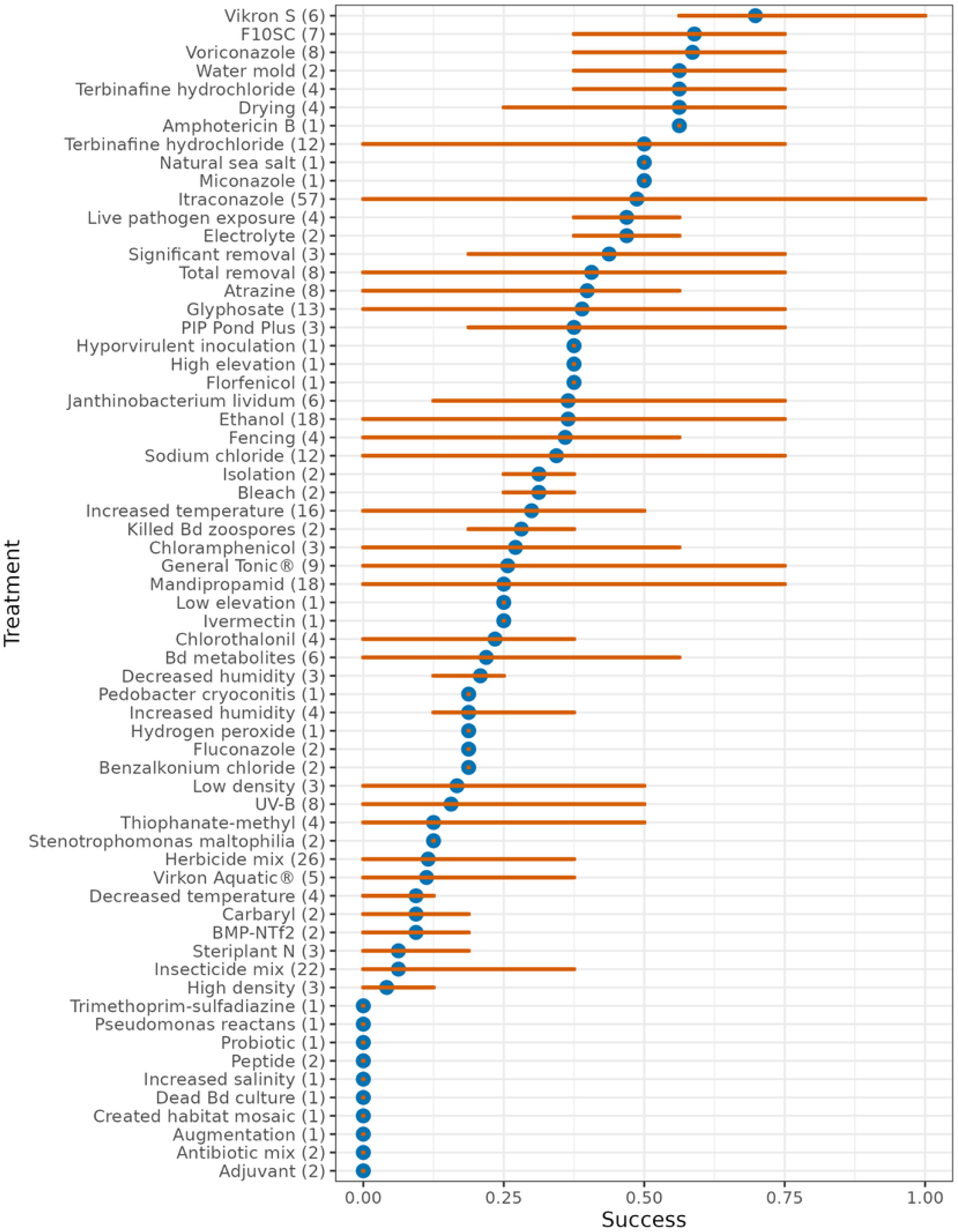
Forest plot showing success of treatments. Blue dots represent mean and orange lines range of success value for that treatment.

**Figure 4.**
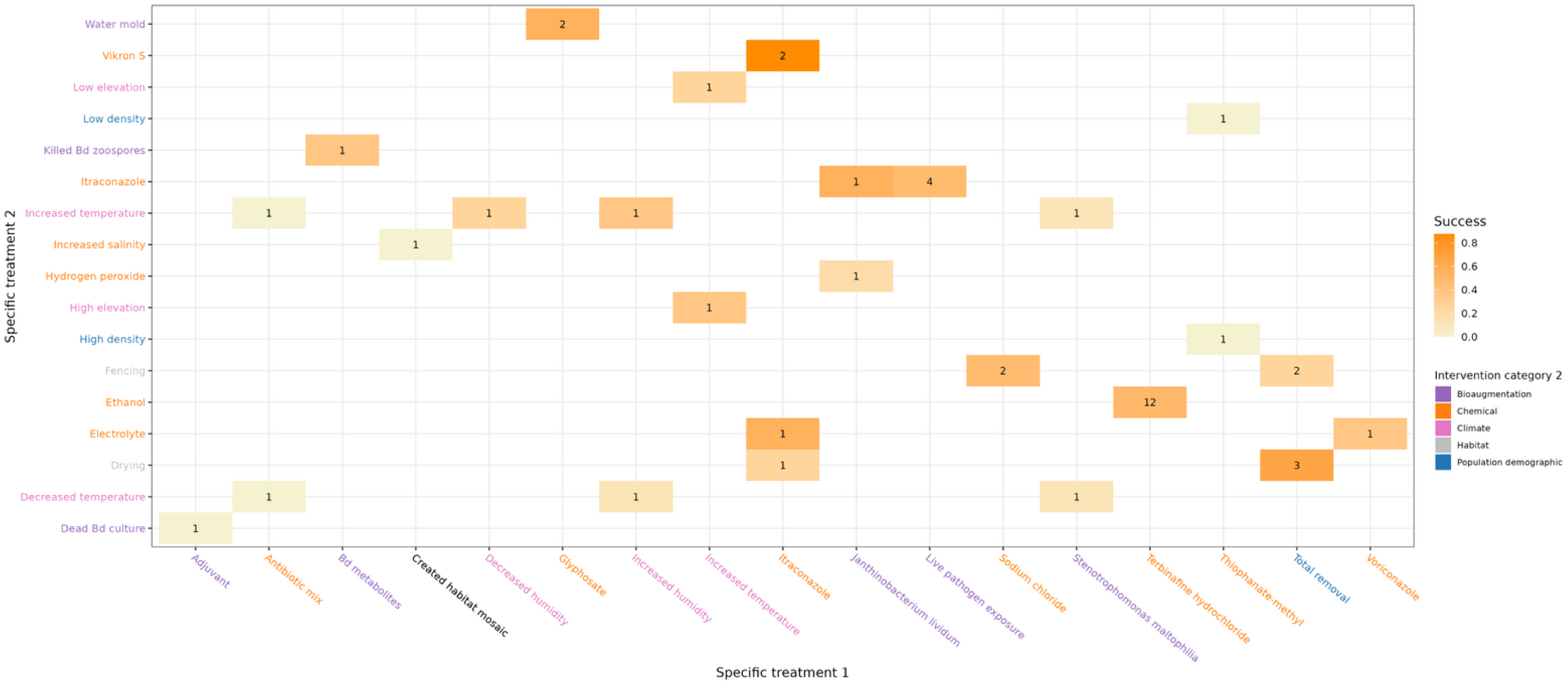
Heat map showing combinations of treatments for all interventions that applied multiple treatments in a single intervention. Labels are coloured by intervention category. Numbers represent count of interventions with that combination of factors. Colour scale represents mean success score for that combination of treatments.

### Interventions across time

Experimental data was published between 2000 and 2023. Many publications did not report year of study but for the 180 that did, experiments took place between 2002 and 2018. A breakdown of the number of studies by category of intervention is shown in Supplementary Figure S2, plotted with time, success and average success for each category per year.

### Patterns of differential success of interventions

Our analyses revealed three main patterns. First, interventions applied to individuals were found to be more successful than interventions applied to the habitat (individual-habitat estimate = −1.11 [−2.07, −0.18]). Second, itraconazole interventions were found to perform better than population demographic interventions (itraconazole-population demographic estimate = −2.17 [−4.33, −0.18]). Third, larvae tended to respond better to treatment than adults (larvae-adult estimate = −0.12 [−2.06, −0.01]). However, the posterior densities of the variables compared to the reference levels (chosen to be the best performing combination: individual, itraconazole, larvae), reveals that the other predictor levels are all skewed towards negative estimates (Figure 5), confirming the best performing combination. Use of other reference levels was also modelled to check robustness and sensitivity, in which individual was shown to perform better than habitat and increased success of itraconazole and larvae was found to be borderline.

**Figure 5.**
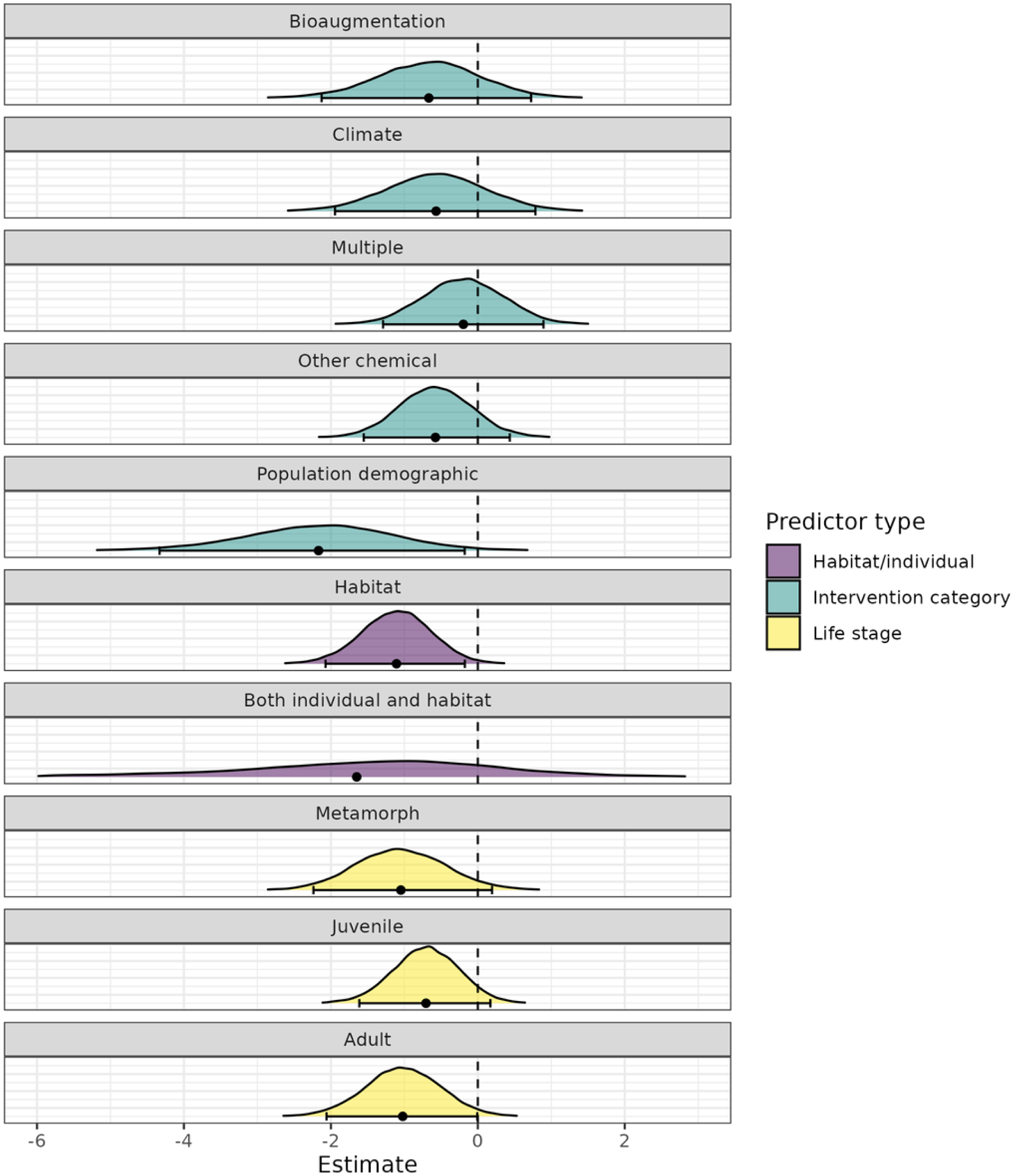
Posterior distributions from best model. Itraconazole, individual and larvae were used as reference levels so each estimate is compared to this combination of predictor factors. Colours correspond to predictor type. Forest plots can be seen along bottom of each distribution, where black dots indicate mean estimate and black lines indicate credible intervals. Habitat interventions statistically perform worse than interventions on individual amphibians. Interventions on adults perform borderline worse than interventions on larvae. Population demographic interventions perform worse than itraconazole interventions.

The probability matrix (Table 1a) shows that population demographic treatments are 90% less likely to be successful than any other treatment. Although it is important to consider that only 13/315 (4%) interventions were population demographic interventions, it should also be noted that 12 of these were on larvae and were classed as individual treatments, which were the best performing factors of that type, with larvae being over 90% more likely to be treated successfully than other life stages (Table 1b), and individual treatments are nearly 99% more likely to be successful than treatments on habitat only (Table 1c). Moderate correlation was found between these three predictors (Figure 6a and Supplementary Figure S3) but not enough to explain these results.

**Figure 6.**
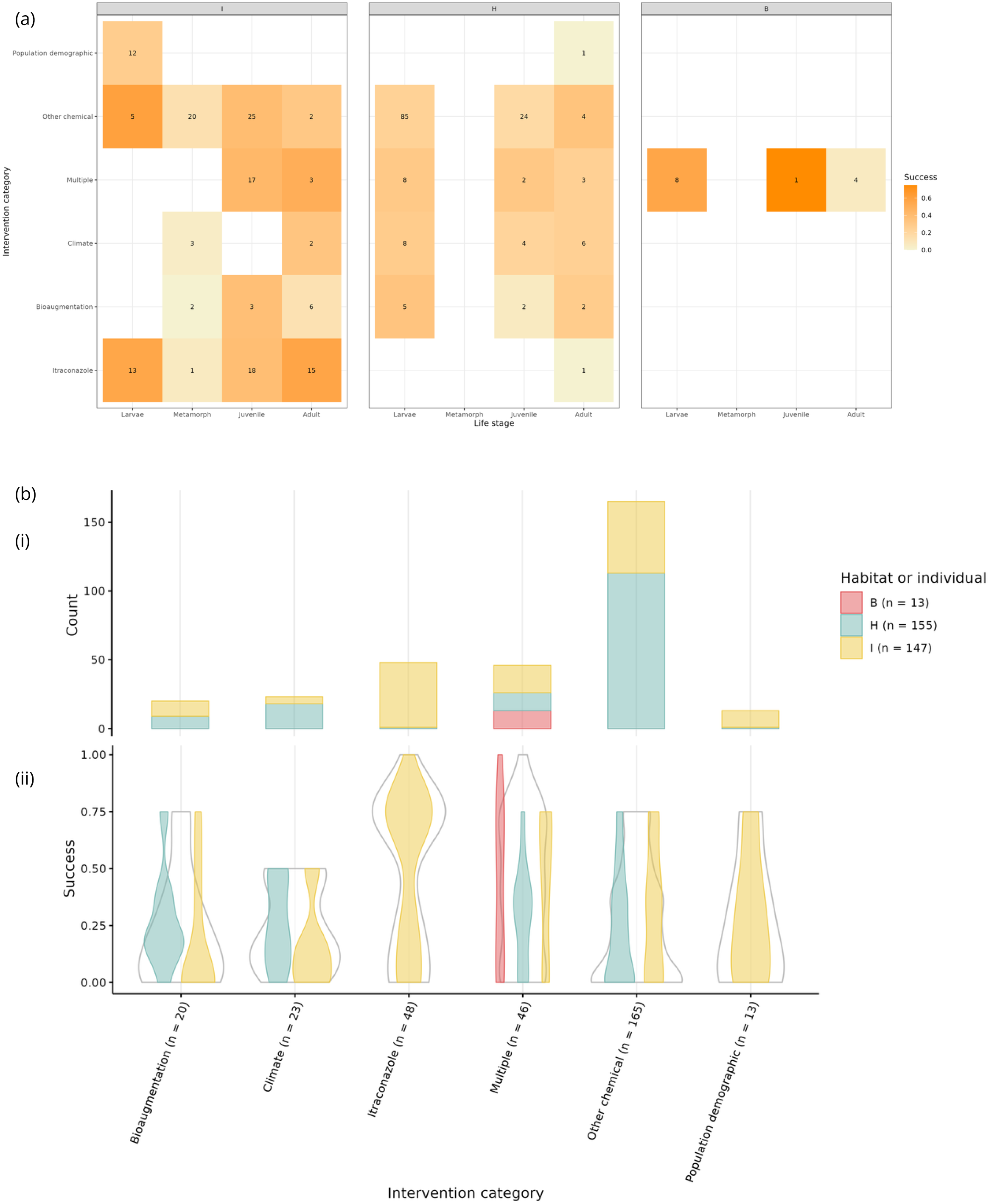
Analysing relationships between pertinent variables. (a) Heat map showing data distribution of intervention category, life stage and habitat. Numbers represent count of interventions with that combination of factors. Colour scale represents mean success score for that combination of factors. (b) Breakdown of success by intervention category and habitat/individual. (i) Stacked bar plot showing number of interventions for each intervention category, broken down by habitat/individual. (ii) Violin plots showing success of each intervention category (grey outline) and split by habitat/individual.

**Table 1.**
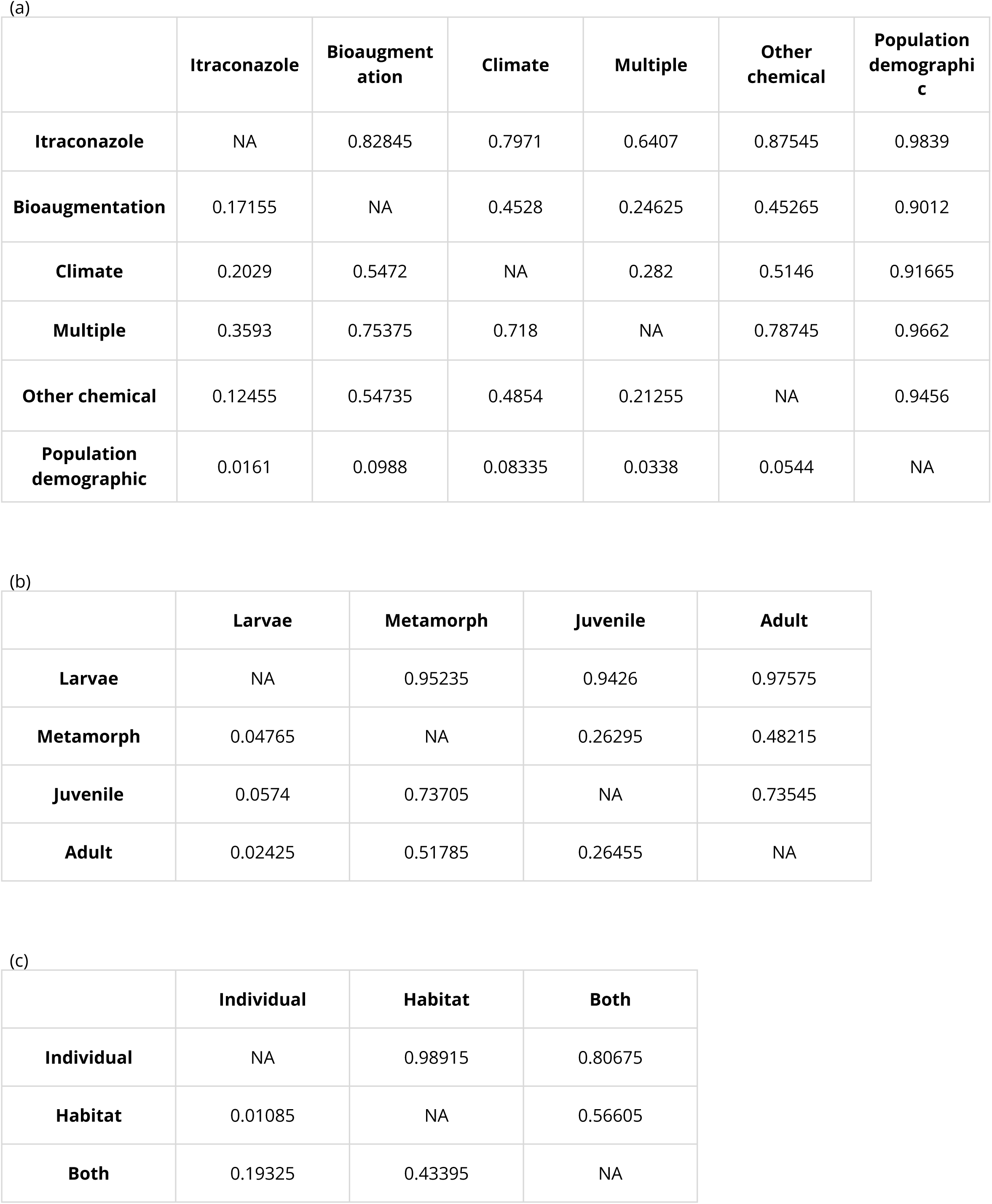
Probability tables based on best model. Individual, itraconazole and larvae were used as reference levels. Cells show probability of higher success of row label compared to column label. (a) comparison of intervention categories; (b) comparison of life stages; (c) comparison of habitat/individual.

|  | **Itraconazole** | **Bioaugmentation** | **Climate** | **Multiple** | **Other chemical** | **Population demographic** |
| --- | --- | --- | --- | --- | --- | --- |
| **Itraconazole** | NA | 0.82845 | 0.7971 | 0.6407 | 0.87545 | 0.9839 |
| **Bioaugmentation** | 0.17155 | NA | 0.4528 | 0.24625 | 0.45265 | 0.9012 |
| **Climate** | 0.2029 | 0.5472 | NA | 0.282 | 0.5146 | 0.91665 |
| **Multiple** | 0.3593 | 0.75375 | 0.718 | NA | 0.78745 | 0.9662 |
| **Other chemical** | 0.12455 | 0.54735 | 0.4854 | 0.21255 | NA | 0.9456 |
| **Population demographic** | 0.0161 | 0.0988 | 0.08335 | 0.0338 | 0.0544 | NA |

(b)
|  | **Larvae** | **Metamorph** | **Juvenile** | **Adult** |
| --- | --- | --- | --- | --- |
| **Larvae** | NA | 0.95235 | 0.9426 | 0.97575 |
| **Metamorph** | 0.04765 | NA | 0.26295 | 0.48215 |
| **Juvenile** | 0.0574 | 0.73705 | NA | 0.73545 |
| **Adult** | 0.02425 | 0.51785 | 0.26455 | NA |

**Table 1.** Probability tables based on best model.
|  | **Individual** | **Habitat** | **Both** |
| --- | --- | --- | --- |
| **Individual** | NA | 0.98915 | 0.80675 |
| **Habitat** | 0.01085 | NA | 0.56605 |
| **Both** | 0.19325 | 0.43395 | NA |

Splitting the data into subsets of interest (larvae-only, adult-only, *in situ*-only, chemical-only, and itraconazole-only) and running separate analyses revealed no further results of interest, except that itraconazole was statistically more successful than other chemical and population demographic in the larva-only model. The *in situ*-only model did not converge, likely due to the low sample size overall and small sample sizes within grouping variables. Dosage was also included in the itraconazole-only model but was not found to influence success.

On further inspection (Figure 6), we see that all but one itraconazole treatments were on individuals, confounding the results showing higher success for itraconazole and individuals and making their effects difficult to untangle. Models run with only a simple binary itraconazole predictor instead of intervention categories showed itraconazole treatments perform better, and models run with only habitat/individual shows individual performs better. However, running the model with itraconazole removed showed no effect of any predictors, including habitat/individual, leading us to conclude that it is itraconazole that is the most pertinent factor in success. However, it should be noted that itraconazole is the only specific treatment with enough separate interventions to allow analysis. Other specific treatments have had to be grouped together, which may mask specific treatment effects (Figure 3).

### Best intervention approaches

There were 62 interventions with a success score of 0.75 or higher (Supplementary Data S2) across 26 publications and 33 species. All received top scores for efficacy and adverse effects, but varied for scalability and longevity, with 5 receiving 1 for longevity and 59 for scalability.

These top interventions contained 8 *in situ* interventions, 46 individually applied interventions (as opposed to 11 habitat and 5 both), spanned 33 different species with 27 larval, 25 juvenile and 10 adult interventions. Of these top scorers, 27 were itraconazole treatments, 18 were non-itraconazole chemical treatments and 13 were interventions with multiple treatment types, leaving only 2 bioaugmentation and 2 population demographic interventions, and no climate treatments with top success scores.

Two interventions received the top score of 1. One is an *ex situ* itraconazole treatment performed on *Alytes obstetricans* (37) and the other was an *in-situ* chemical intervention that used a combination of 1 mg/l itraconazole applied to the individual amphibian and 0.1% Virkon S applied to the environment, performed on *Alytes muletensis* (22). However, it should be noted that neither received the top usability score, with lower confidence intervals of 0.995 for Bosch et al. (22) and 0.999 for Tobler and Schmidt (37).

## METHODS

### Study selection

Studies were selected if they met the following criteria: peer-reviewed publications on experimental trials applying intervention methods *in vivo* individuals, their habitat, or in combination, and which were tested against *Bd*-positive control groups or *Bd* infection load/prevalence of the same trial group prior to intervention. The terms “chytrid fungus,” “chytridiomycosis,” and “*Batrachochytrium dendrobatidis*” were searched in combination with “mitigation,” “treatment,” and “intervention” on both the Web of Science and Google Scholar. Results were ordered by relevance, and for each combination of terms, the abstracts of the first 300 results were reviewed to determine whether they met our criteria.

Studies that focused on population monitoring without implementing any treatment were excluded. This included research observing the impact of *Bd* under varying climatic or geographic conditions in natural settings, unless those variations involved intentional habitat manipulations that alter natural conditions.

### Data extraction

The following data was collected and from each study trial: (1) focal species, (2) the origin of population, (3) study location, (4) year of treatment, (5) *in situ* or *ex situ*, (6) the number of individuals in the treatment group, (7) life stage of treated population, (8) intervention category(s) (Supplementary Table S1a), (9) specific treatment(s) used, (10) duration of treatment application, (11) monitoring for more than six months post-treatment (yes or no), (12) targeted application of the treatment on the individual, habitat or both, (13) therapeutic or prophylactic application of the treatment, and (14) dosage or temperature of treatments.

Three different scoring systems were developed to assess the efficacy of intervention for each treatment trial (Supplementary Table S1b), where efficacy was taken to be the difference between the control or pre-treatment group and the treatment group six months (or the next nearest measurement after six months) after treatment, or the maximal difference if this was unavailable. The ‘infection matrix’ assessed the log_10_-transformed difference in zoospore load. The ‘prevalence matrix’ compared the proportion of individuals infected with *Bd* in the control or pre-treated group to the proportion cleared of infection in the treatment group. The ‘survivability matrix’ assessed the proportionate survivability between groups. ‘Infection matrix,’ ‘prevalence matrix,’ and ‘survivability matrix’ were prioritized respectively. The ranking order was determined by two factors: frequency of use across trials, and reduction of ambiguity in both presentation and gathering of results.

A ‘trend in efficacy score’ was recorded for any entry with long-term monitoring in which there were additional data points after the point from which the efficacy score was assigned. It was assigned one of ‘no change’, ‘increase’, or ‘decrease’ in the score, determined by a comparison of efficacy between the point at which the efficacy score was assigned and the efficacy score evaluated after the six-month monitoring period. If there were no additional data points or if the study did not have any long-term monitoring, it was assigned N/A. This value was then reduced to a constructed data field called ‘longevity’, which assigned a 1 for any trial with no change or an increase in efficacy after long-term monitoring, and a 0 otherwise (hence, trials with no monitoring were given the same score as trials with a decrease in efficacy over time). ‘Adverse effects’ were scored based on whether the application of the treatment had no negative impacts (zero), caused distress (one), caused death to 25-50% of the treatment group (two), or caused the death of more than 50% of the treatment group (three). Each entry was also assessed on whether the intervention was scalable or not. ‘Scalability’ was encoded as a binary (yes or no) value. It was determined by whether the intervention type is suitable for a large distribution without unreasonable requirements of additional labour and cost (i.e., requiring repetitive, long-term treatment applications but easily introduced into the daily tasks of personnel) and can be implemented both *ex situ* and *in situ*. Lastly, each entry was assigned a score of zero to three based on ‘usability’ (Supplementary Table S1c), where zero is the top score. This quantifies whether the methods, data collected, experimental setup, and important information were presented in the paper to fulfil all the data fields later used in the analysis.

### Calculating intervention success

Using the methodology described in (38), we used the data extracted above to calculate a success value based on the efficacy of the intervention, accounting for adverse effects, and scaled by the longevity and scalability of the intervention, with these designations as described above. The weights of these scaling factors were chosen to give equal weight to scalability and longevity at 0.25 respectively, and 0.5 for the baseline weight (the maximum score that can be achieved if neither the scalability nor longevity criteria are met). Success was calculated based on this configuration of variables (Supplementary Figure 1a, b). Any intervention with a usability of 3 was removed and weights to be used in the meta regression (Supplementary Figure 1c) were calculated following (38) where usability was scaled so that those interventions with the lowest usability score of 2 were given 60% weight, 80% to those with a score of 1 and 100% to those with the best score of 0 (Supplementary Figure 1d). These weightings were multiplied by a modified sampled size measure, scaled between 0 and 1 using a sigmoid function on the log10 of the sample size and a cap of 600, whereby any intervention with over 600 samples was given a scaled sampled size of 1 (Supplementary Figure 1d). Any unknown sample size was given a raw sample size value of 1, which was then used in the scaling process described.

### Publication bias

Studies with positive results tend to be published more readily than studies with negative results. Therefore, this publication bias must be considered when conducting assessments such as this. Although 33% (101/313) received a success score of 0, we do not know how many more studies with negative results have not been published, therefore we must attempt to account for these in our analysis. However, since our dataset is purposefully highly heterogeneous (with different interventions, methodologies, species, etc.), it is very difficult to disentangle publication bias from these effects. The funnel plot is therefore more like a scatter plot because of the heterogeneity and so is not useful for assessing publication bias. Since our study compares metrics across studies rather than looking at the overall success score, we are required to make the assumption that negative results are lacking across each variable type equally, and focus on the meta-regression to try and disentangle these effects. Equally, certain subsets of variables may be more likely to be published, such as certain species, certain intervention types etc. and we were only able to use English language studies in our analysis.

Further, since our effect size measure, success, is beta-distributed, standard error naturally decreases towards success values of 0 and 1. All of these may impact the results, so we should use caution when extrapolating the results from this dataset to more generalised settings.

### Statistical analyses

All models were conducted in brms (39) and run for 10^4^ iterations, including 5×10^3^ burn-in, and using the default uninformative priors (half student t distribution (df=3, location=0, scale=2.5) for random effects and improper uniform distribution across the real numbers for fixed effects). Leave-one-out cross validation was used to evaluate each model and compare models with each other by assessing the difference in ELPD and its associated standard error, also using the brms package (39).

We explored the grouping variables (taxa (as genus), publication and efficacy matrix) by running random effects models. Publication and efficacy matrix were strongly associated with success, but not genus. However, as publication and efficacy matrix are also strongly correlated with each other and the difference between the models (including the model with both) was negligible within error, our final models control only for publication.

We ran models with the following variables: *in situ/ex situ*, life-stage, intervention category, habitat or individual, multiple or single treatment categories, therapeutic vs prophylactic, tropical or temperate amphibian species, terrestrial or aquatic amphibian habitat, mean snout-vent-length, minimum clutch size, and amphibian taxa. Correlation analysis was performed to test for collinearities, and bivariate models were run to be used in conjunction with multivariate models for predictor selection and exploring the results.

## DISCUSSION

Our study provides the first global-scale quantitative assessment of differential successes among the wide range of interventions applied to mitigate the eroding impacts of *Bd* on Anthropocene amphibian biodiversity. Our evidence reveals two main phenomena—one biological and one geoeconomic. Biologically, the majority of interventions offer unreliable degrees of success, whereas the implementation of itraconazole-based chemical interventions applied to individual tadpoles have the most consistent rate of success. Geoeconomically, we observed a dominant tendency for programs implementing interventions to concentrate disproportionately across the Global North and its native species, whereas the Global South has been drastically lagging behind.

Although chemical treatment with itraconazole has been shown to be highly effective as a treatment in laboratory settings, its effectiveness in the wild remains to be addressed, given that only 15% (7/48), or 19% (11/57) including combinations, of itraconazole interventions were tested in the field. This is a trend across all interventions, with 90% of all trials being performed *ex situ—*an unsurprising proportion due to the increased complexity of performing *in situ* experiments. Given this bias, our results are naturally dominated by the patterns found in *ex situ* interventions, so. while an important first step, *ex situ* results and trends leave many unknowns when it comes to real-world implementation. Therefore, despite *ex situ* success, whether the same type of intervention applied *in situ* will have the same successful effects remains unknown. Therefore the observed success of the itraconazole-based interventions must be considered within the context of its application, despite our attempts to mitigate this with our inclusion of additional criteria into the success calculation.

Itraconazole is an anti-fungal chemical and using it in natural environments may have unknown negative consequences. Fungicides are commonly used in human and animal medical treatments, as well as to protect crops against fungal diseases, and have been known to have negative side effects such as increasing pathogen resistance, harming non-target species and contaminating the environment (40,41). Therefore, widespread use of fungicides to treat *Bd in situ*, although a seemingly promising solution in terms of efficacy, should be approached with caution. In addition, itraconazole has been applied for the past three decades as a treatment for human fungal diseases (42). Therefore, before being trialled as a *Bd* treatment, it had already undergone substantial research, which means it did not require preliminary work in testing its effects as an anti-fungal. This may explain its earlier successes and its predominance in treatment types as an initially more promising solution, in comparison to interventions like bioaugmentation, which are relatively novel. Therefore, despite the lower success rate of the non-chemical options, these could still provide a promising avenue of treatment, particularly since most chemical treatments are inspired by human and agriculture disease mitigation, which may be inappropriate in natural environments.

In general, however, the observation that itraconazole-based interventions excel in success reinforces the empirical evidence that pharmacological approaches are currently superior when treating the outbreak of infectious diseases. This result is not surprising as chemical interventions are universally accepted methods for treatment of diseases. Importantly, however, our results also appear to resemble the trends previously observed in human medical science—for example, the geographic structure of the so-called “vaccine apartheid”, which shaped the geopolitical and geoeconomic distribution of unequal access to vaccines during the COVID-19 pandemic (43–45). Increased funding and higher capacity in the Global North frequently means that countries in the Global South, which are often impacted more severely by disease, are unable to perform the research they need themselves. We can also see here that not only are the majority of trials being tested the Global North, but the species used are also mostly native to these countries, despite the fact that the hotspots of amphibian extinctions (overall, not specifically the declines attributed to *Bd*), and of amphibian biodiversity are disproportionately concentrated in low/middle-income countries from tropical regions (12,46). This raises a further question of whether we need to consider active discussions to re-think a more globally cooperative agenda to meet the challenges of understanding the strengths and limitations of interventions, and to ultimately mitigate the devastating impacts of the Bd infectious disease on amphibian persistence.

## Author contributions

Conceived the study: LEBG and DPD; Data collation: VCT; Data analyses: LEBG; Manuscript writing: LEBG, VCT and DPD.

## Data availability

The dataset used in this study is available at Figshare

## Code availability

Code is available at

