## Supplementary Materials for "Global assessment of interventions for mitigation of amphibian fungal disease is dominated by geoeconomic trends and antifungal success"

**Code.** All code used in the analysis.
Github link: https://github.com/lebgoodyear/2025_bd_interventions

**Data**
**S1.** Full collated dataset and metadata in xlsx format.
**S2.** Full dataset used in analysis in csv format.
**S3**. Dataset in csv format containing 62 interventions with a success score of 0.75 or higher.
DOI for all datasets: 10.6084/m9.figshare.29416430

**Figures S1-S3.**

**Table S1.**

**Supplementary references.**

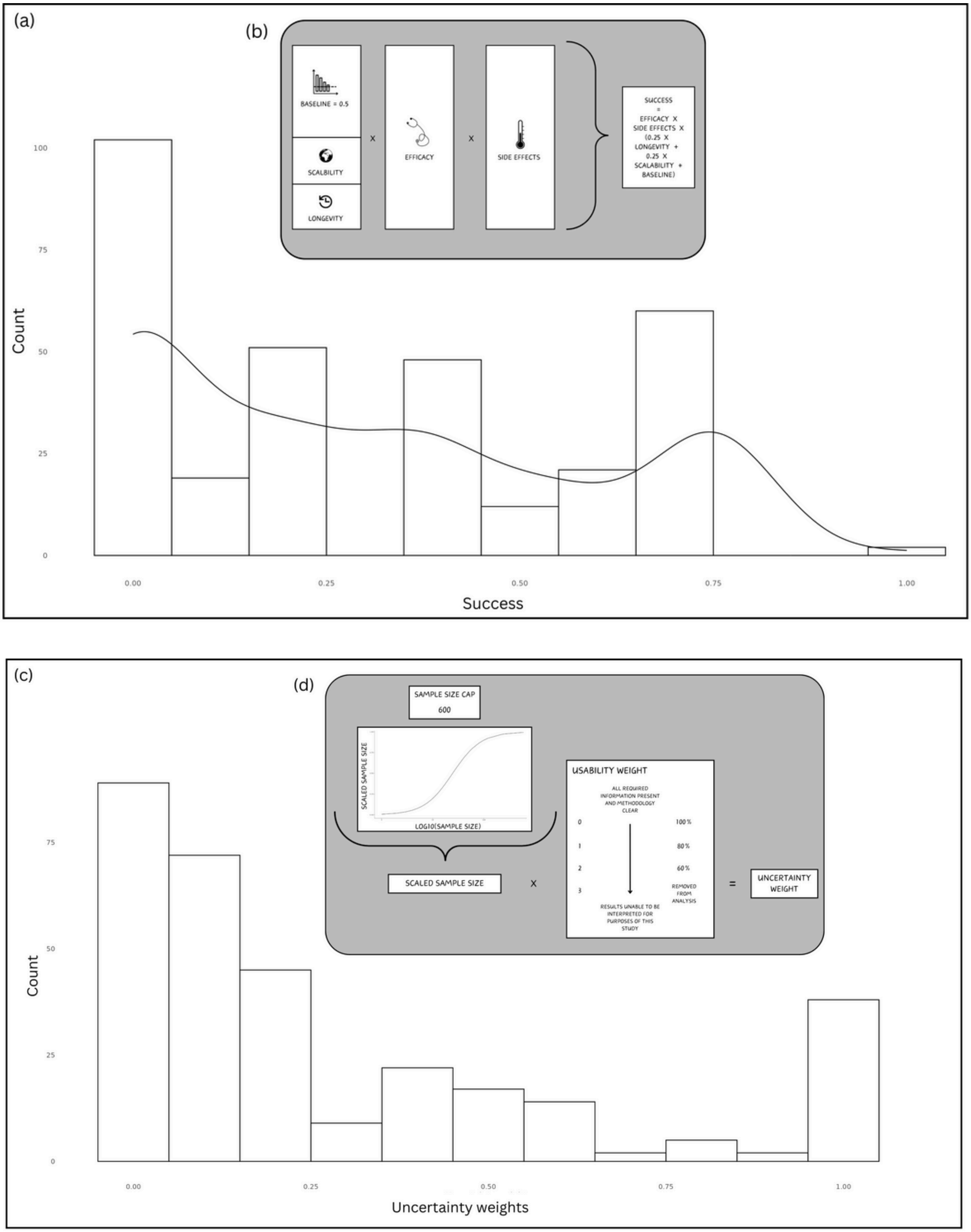


**Figure S1.** Success and uncertainty weights distributions. (a) Success calculation breakdown. (b) Distribution of success in dataset. (c) Uncertainty calculation breakdown. (d) Distribution of uncertainty weights in datasets.

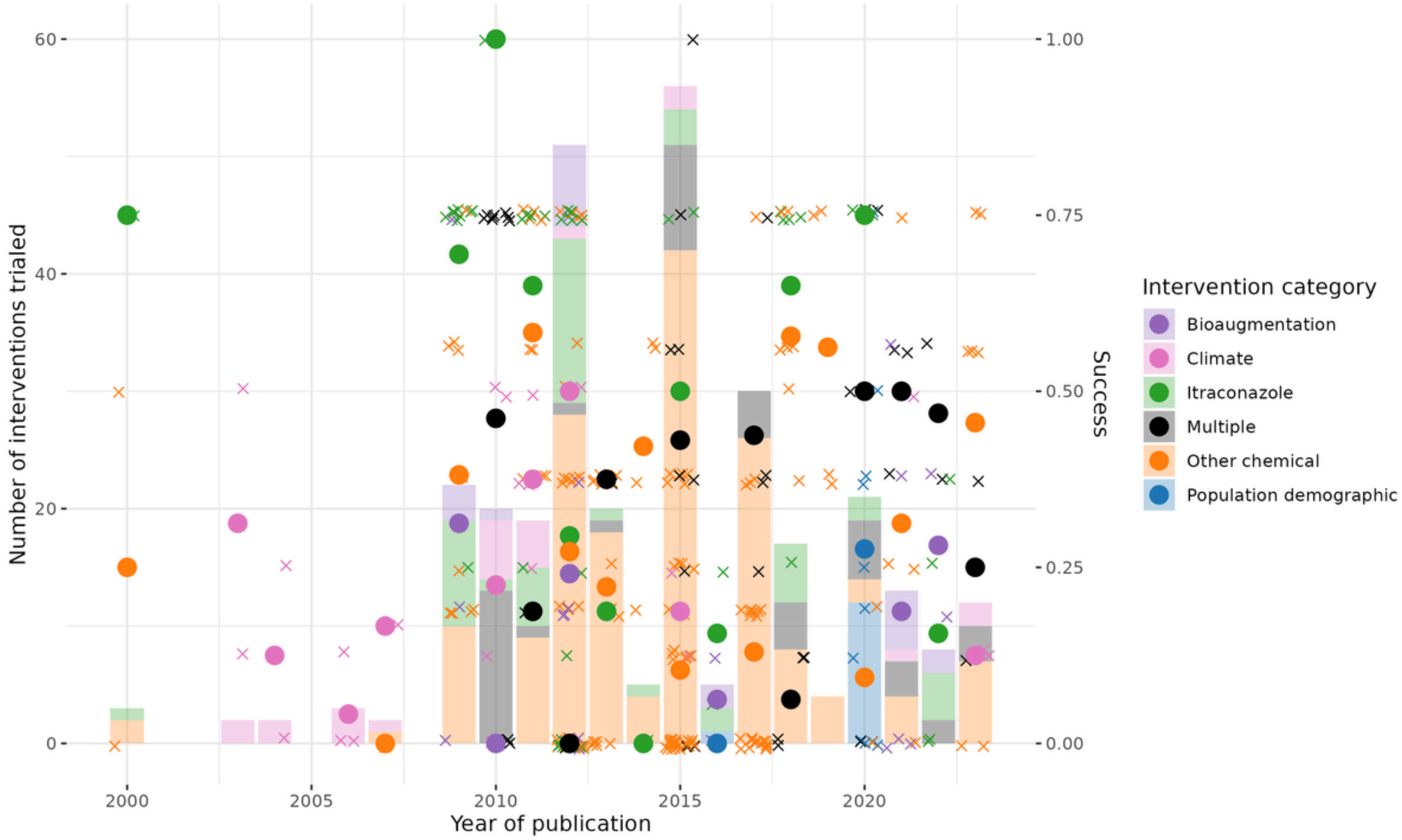


**Figure S2.** Plot showing numbers of interventions trialled per year (left y-axis), broken down by intervention category. Right y-axis shows success score for each intervention (crosses) and the average success score per intervention per year (filled circles).

(a)

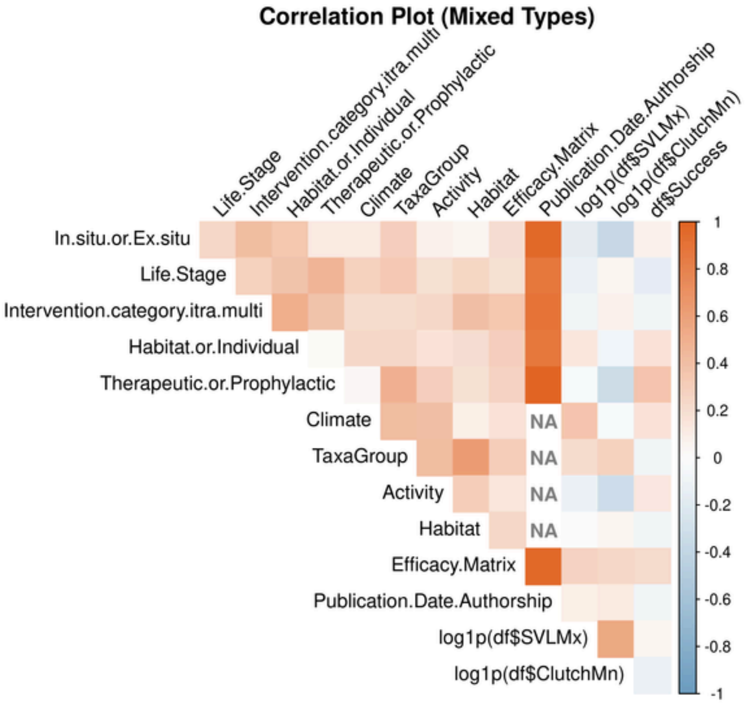


(b)

| | In.situ.or.Ex.situ | Life.Stage | Intervention.category.i | Habitat.or.Individual | Therapeutic.or.Prophy | Climate | TaxaGroup | Activity | Habitat | Efficacy.Matrix | Publication.Date.Autho | log1p(df$SVLMx) | log1p(df$ClutchMn) | df$Success |
|---|---|---|---|---|---|---|---|---|---|---|---|---|---|---|
| In.situ.or.Ex.situ | 1.000 | 0.245 | 0.412 | 0.356 | 0.122 | 0.129 | 0.296 | 0.089 | 0.057 | 0.202 | 0.965 | -0.165 | -0.365 | 0.079 |
| Life.Stage | NA | 1.000 | 0.283 | 0.374 | 0.461 | 0.287 | 0.334 | 0.198 | 0.256 | 0.199 | 0.872 | -0.129 | 0.050 | -0.153 |
| Intervention.category.itra.multi | NA | NA | 1.000 | 0.515 | 0.375 | 0.215 | 0.223 | 0.242 | 0.397 | 0.359 | 0.900 | -0.083 | 0.077 | -0.085 |
| Habitat.or.Individual | NA | NA | NA | 1.000 | 0.021 | 0.232 | 0.246 | 0.160 | 0.210 | 0.292 | 0.872 | 0.148 | -0.072 | 0.168 |
| Therapeutic.or.Prophylactic | NA | NA | NA | NA | 1.000 | 0.039 | 0.494 | 0.297 | 0.185 | 0.274 | 0.993 | -0.031 | -0.332 | 0.362 |
| Climate | NA | NA | NA | NA | NA | 1.000 | 0.428 | 0.406 | 0.099 | 0.179 | NA | 0.369 | -0.040 | 0.167 |
| TaxaGroup | NA | NA | NA | NA | NA | NA | 1.000 | 0.430 | 0.628 | 0.320 | NA | 0.215 | 0.290 | -0.085 |
| Activity | NA | NA | NA | NA | NA | NA | NA | 1.000 | 0.306 | 0.155 | NA | -0.110 | -0.313 | 0.135 |
| Habitat | NA | NA | NA | NA | NA | NA | NA | NA | 1.000 | 0.249 | NA | -0.021 | 0.055 | -0.085 |
| Efficacy.Matrix | NA | NA | NA | NA | NA | NA | NA | NA | NA | 1.000 | 0.967 | 0.274 | 0.243 | 0.224 |
| Publication.Date.Authorship | NA | NA | NA | NA | NA | NA | NA | NA | NA | NA | 1.000 | 0.092 | 0.129 | -0.093 |
| log1p(df$SVLMx) | NA | NA | NA | NA | NA | NA | NA | NA | NA | NA | NA | 1.000 | 0.560 | 0.053 |
| log1p(df$ClutchMn) | NA | NA | NA | NA | NA | NA | NA | NA | NA | NA | NA | NA | 1.000 | -0.129 |
| df$Success | NA | NA | NA | NA | NA | NA | NA | NA | NA | NA | NA | NA | NA | 1.000 |

**Figure S3.** Correlation matrix for all variables. (a) Heat map with correlation strengths. (b) Matrix with actual correlation values (Pearson Correlation Coefficient, Polyserial Point Correlation or Cramer's V). Matrix is symmetric and repeated values have been set as NA for ease of viewing.

(a)

| Intervention Category | Description | General Treatment(s) |
|---|---|---|
| Chemical Application | Any treatment to the test subjects or their habitat that requires adding chemical drug treatments or any other *Bd*-inhibiting substance that is abiotic (Bosch *et al*. 2015; Drawert *et al*. 2017; Pessier 2014). Included in this is the cleaning of a habitat that would require draining for the purpose of cleaning it with a chemical compound prior to the re-introduction of a population. | • Antifungal<br>• Antibiotic<br>• Antibacterial<br>• Antiparasitic<br>• Sodium<br>• Disinfectant<br>• Herbicide<br>• Insecticide |
| Habitat Manipulation | Any change to the habitat of the host population that includes only physical manipulation (Scheele *et al*. 2014). This manipulation can inadvertently result in a change of temperature or features in the environment. Creation of new features or new habitat mosaics, such as basking/bathing rocks, water features, blocking of cool water reservoirs, would be included in this category (Klop-Tocker *et al*. 2021). Cleaning of a habitat with either a disinfectant or antifungal would not be considered in this category, but if a location is drained and dried with no application of chemical compound then it would be included here. | • Fencing<br>• Drying<br>• Creation of habitat<br>• Removal/Addition of environmental features (e.g., vegetation, bathing stones, cold water refuges) |
| Climatic Manipulation | Any manipulation of climatic factors that results in either the condition of the habitat to be outside of the optimal range for *Bd* growth, or any treatments applied to subjects that result in experiencing conditions outside of that same range (e.g., temperature changes outside of a 15-25°C) (Brannelly *et al*. 2015; Gass *et al*. 2023; Greenspan *et al*. 2017). | • Temperature<br>• Humidity<br>• Radiation |
| Population Demographic Adjustment | Within an extant population, any adjustments to the composition of that population's demographic (Bosch *et al*. 2020; Dobson & Carper 1996; Fernández-Loras, Boyero & Bosch 2020; Scheele *et al*. 2021). | • Augmentation/Removal of overall population, age or gender composition<br>• Density<br>• Reintroductions |
| Bioaugmentation | Introduction of bacteria or any other biotic additions to either habitat or directly to the subjects to stimulate immunological response (Bletz *et al*. 2013; Harris *et al*. 2006, 2009a, 2009b; Muletz *et al*. 2012; Stice & Briggs 2010). | • Microbiota<br>• Live/Dead pathogen exposure<br>• Commensal bacteria<br>• Metabolite<br>• Probiotic<br>• Peptide<br>• Inoculation<br>• Adjuvant |
| Translocation | Any movement of subjects or population to a location outside of species subject's historic range (Scheele *et al*. 2021). | • Introduction |

(b)

| Efficacy Score | Infection | Prevalence | Survivability |
|---|---|---|---|
| **0** | No change or an increase in infection load. | No change or increase in the presence of infection | No change or decrease in survivability. |
| **1** | Differences in units of measurement between control/pre-treatment and the treatment groups was between 0 and 0.5 (log10). | Less than a 30% reduction in the presence of infections, or only 30% of the population tested negative. | Survivability was less than 1.5 times higher than control, or 0-40% of the population survived. |
| **2** | Differences in units of measurement between control/pre-treatment and the treatment groups was between 0 .5 and 1.5 (log10). | Between a 30-70% reduction in the presence of infections, or 30-70% of the population tested negative. | Survivability was 1.5-3 times higher than control, or 40-70% of the population survived. |
| **3** | Differences in units of measurement between control/pre-treatment and the treatment groups was more than 1.5, but not a total elimination of zoospores (log10). | More than a 70% reduction in the presence of infections, or more than 70% of the population tested negative. | Survivability was more than 3 times higher than control, or 70-99% of the population survived. |
| **4** | The treatment group completely eradicated the infection compared to the control/pre-treatment group. Zero *Bd* zoospores (log10) were detected. | No presence of infection in the population. All individuals in the treatment group tested negative. | 100% survivability. |
| N/A- Results were not interpretable for the purposes of this study. | | | |

(c)

| Usability Score | Definitions |
|---|---|
| **0** | All information was present, and the methodology was clear. The experimental setup was clear and well-understood. |
| **1** | Up to two salient data fields were not present in the paper or supporting information and/or the methodology and the experimental design was not clear and lead to inability to divide information into data fields. |
| **2** | More than two salient data fields were not filled because of the lack of information in the paper or supporting documents and/or the experimental set-up was found to be convoluted for the purposes of this study (e.g., testing more than two interventions within experimental groups). |
| **3** | The paper is not suitable. |

**Table S1.** Tables of definitions used for sorting relevant information and data collection metrics. (a) Categorisation of the interventions used in mitigations with description and treatment examples for each category. (b) Efficacy scoring system with description of each of the three scoring matrices and their scoring rubric used to assign an efficacy score to each treatment trials. (c) Usability definitions to assign a score to each treatment trial to determine suitability for the intents and purposes of this research. Salient fields of data are the predictor variables used in the analysis.
